# Modelling 3D tumour microenvironment *in vivo:* a tool to predict cancer fate

**DOI:** 10.1101/2021.09.23.461461

**Authors:** J. Marines, F. Lorenzini, K. Kissa, L. Fontenille

## Abstract

Recently, many studies demonstrated the fundamental role of tumour microenvironment (TME) in cancer progression. Here, we describe a state-of-the-art method to visualize in 3D the behaviour of tumours in zebrafish embryos. We highlight two major actors of TME, macrophages and vessels. This valuable tool is transposable to Patients Derived Xenograft imaging in order to predict the fate of malignant tumours according to the dynamics of their TME.

---

The TME characterisation has become over the years a major topic in the understanding of tissue tumorigenesis and the development of treatments. The tumour niche is a very dynamic region where different cell types coexist and interact with cancer cells and condition their fate^1^. Among them, vessels play an important role as a pathway for cancer dissemination^2^ and, at the same time, for oxygen and nutrient supply ^3^. Similarly, the recruitment and infiltration of several types of immune cells in the tumour such as macrophages, neutrophils and lymphocytes have been described as having a direct impact on cancer progression ^4^. Finally, the composition of the tumour niche is nowadays used as a mean of diagnosis and indicator of good or bad prognosis ^5 6^. Therefore, the 3D dynamics visualization of the tumour microenvironment appears as a predictive tool of cancer cell behaviour in terms of intravasation, invasiveness and metastasis. To this end, extensive efforts are currently made to explore 3D reconstruction of human cancer biopsies^7 8^. However, to study in detail the molecular and cellular mechanisms of cancer progression, innovative methods based on *in vivo* models are essentials.

The use of *in vitro* immortalized cell lines has been the gold standard for many years. However 2D and 3D culture models remain limited by the number of co-cultured cell types which do not sufficiently account for the complexity of the TME ^9^. Thus, *in vivo* models are crucial to reliably reproduce the interactions between different cell types. In recent years, zebrafish embryos have emerged as a relevant tool in oncology. The simple manipulation, the low cost of rearing and the wide range of genetic manipulations make it one of the most pertinent tools for cancer *in vivo* studies. In addition, its transparency allows to image and visualize in real time conserved cellular processes involved in tumour dissemination as well as interactions between cancer cells and their microenvironment ^10^. Xenograft of human cancer cells in zebrafish embryos have been widely used to model different types of cancer ^11^. Indeed, due to the immaturity of its adaptive immune system at the embryonic stage, rejection after transplantation of human cancer cells is prevented ^12^. Over the last decades, several teams have characterized the types of interactions between xenografted human cancer cells and macrophages in the zebrafish embryo ^13 14^. Furthermore, the interaction between cancer cells and vessels has been intensively studied. Processes such as angiogenesis and vessel co-option reported in patients are conserved after xenografting glioblastoma and melanoma into the zebrafish embryo ^15 16^. The feasibility of simultaneous imaging of three different populations in the yolk sac using two zebrafish reporter lines has been shown ^17^. However, 3D and 4D visualization of macrophages and surrounding vessels in the TME has never been modelled.

Here we report for the first time a cutting-edge methodology to visualize in 3D live imaging the interaction between tumour microenvironment and cancer cells to study cancer fate. To validate and demonstrate the reproducibility of our approach, two different 3D cancer models were established. In order to follow and predict the tumour’s fate, fluorescent human glioblastoma (U87) and human melanoma (A375) cell lines were engineered and transplanted into the midbrain and swim bladder of early-stage zebrafish embryos. Using two zebrafish transgenic reporter lines, *Tg(kdrl:gfp)* and *Tg(mfap4:mcherry)*, we were able to simultaneously study the behaviour of macrophages and vessels in contact with human cancer cells and the fate of the latter.

The general method consists of five steps synthesized in the Fig. 1A. The workflow includes the generation of the stable fluorescent blue lines, the embryos and cells preparation, the xenotransplantation procedure, the image and video acquisition followed by analysis using Fiji and Imaris software. One day after transplantation, we visualised the recruitment of macrophages around glioblastoma (Fig. 1 C) and melanoma tumour (Fig 1 H). Using the Imaris Software, we reconstructed in 3D, the tumour mass and the TME, including the macrophage population and vasculature (Fig 1 F, K). From the 2D images and Z-stack we observed that recruited macrophages interact with cancer cells. However, only 3D visualisation allows to characterize the physical orientation of macrophages around the tumour and their infiltration inside the tumour mass (Sup. Video 1 and 2). In addition, thanks to time-lapse images, the dynamics of macrophages was monitored in 4D (Sup. Video 3 and 4). Long-contact between macrophages and glioblastoma cancer cells are visualised, suggesting an intense communication between the two types of cells (Sup. Video 3). Recently the pro-tumoral role of macrophages in zebrafish glioblastoma and melanoma models has been demonstrated ^13 18^. Quantifying the number of macrophages and following their dynamics in time and space will unravel the steps leading to the evolution of the tumour and the appearance of cancerous metastases.

**Figure 1.**
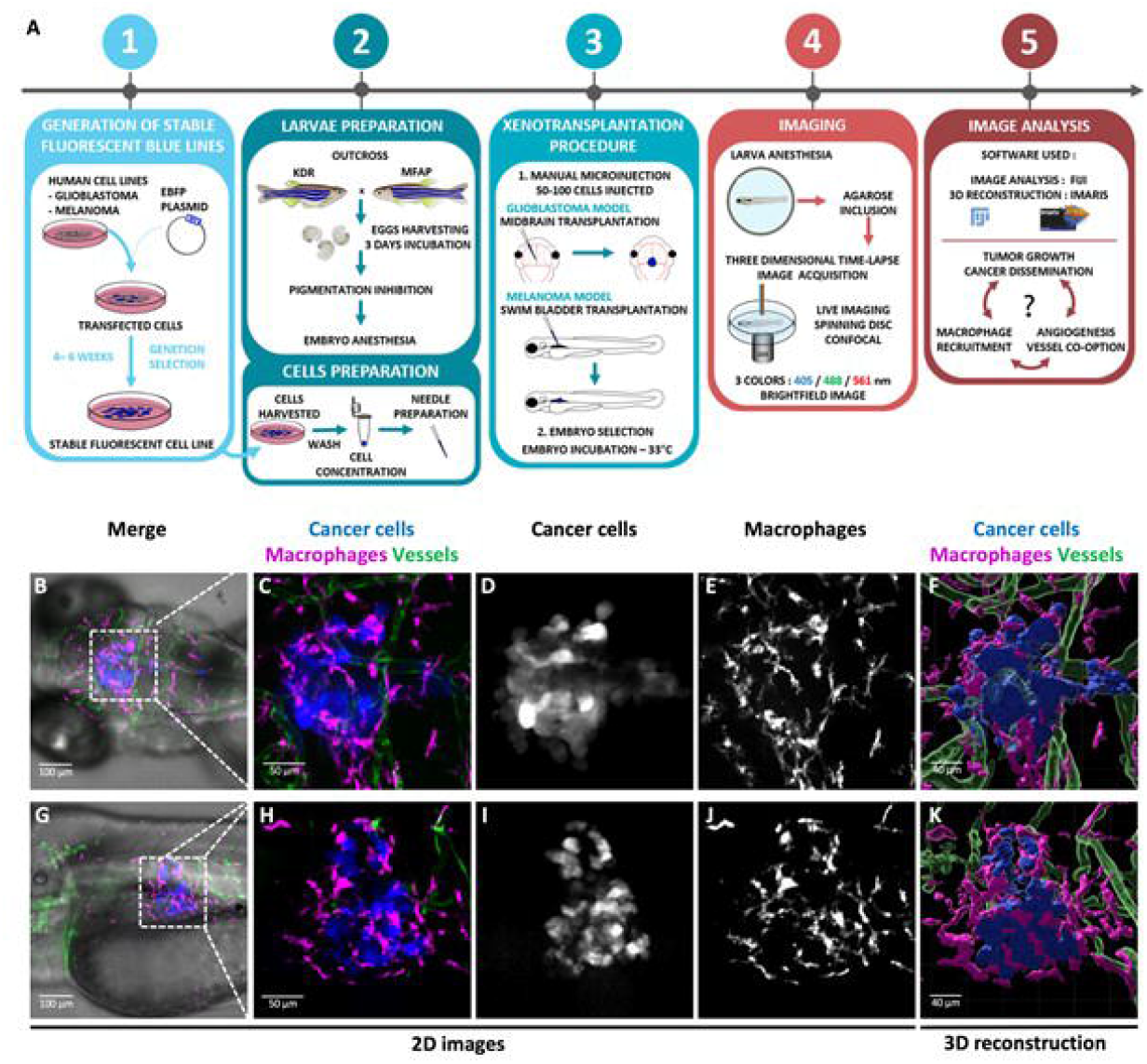
Experimental workflow and 3D glioblastoma and melanoma TME reconstruction. **A**. The method is composed of five successive steps. Blue glioblastoma (GBM) and melanoma fluorescent human cancer cell lines, (U87 and A375 respectively) were firstly generated (1). For this purpose, GBM cells and melanoma cells were transfected with a EBFP2 plasmid and selected during several weeks using Geneticin (1). Then, cells and embryos were prepared prior to transplantation (2). GBM cells were transplanted in the midbrain and melanoma cells in the swim-bladder of 3dpf embryos (3). Imaging and time-lapse were performed at 24h or 48h post transplantation (4). Image analysis was achieved using Fiji and Imaris software (5). **B-K**. Each cell population is highlighted with a coded color; cancer cells in blue or white, macrophages in magenta or white, and vessels in green. **B-E**. Visualization of GBM TME in the midbrain of zebrafish embryo at 4dpf and 24h post transplantation. Images represent a Z projection of 67 slices. **G-J**. Visualization of Melanoma TME in the swim-bladder of zebrafish embryo at 4dpf and 24h post transplantation. Images represent a Z projection of 74 slices. **F**,**K**. 3D reconstruction of tumors infiltrated by macrophages and vessels. **E**,**J**. A High number of macrophages are recruited all around the tumor sites.

We also investigated tumour vasculature evolution, a crucial player in TME. One or two days after cancer cell transplantation, we observed angiogenesis and vessel co-option phenomena. We observed dramatic changes in vessel morphology upon contact with human glioblastoma and melanoma cells. Proximity to human glioblastoma cells directly influences vessel dilation and shape (Fig 2 B,C,G,H and Sup Video 4). The 3D reconstructions highlight the abnormal tortuosity of the vessels (Fig 2 D) and their infiltration within (Fig 2 I and Sup Video 4) the tumour. In the melanoma model, the cancer cells induce neoangiogenesis (Fig 2 L, M). The 3D reconstruction allowed visualization of infiltrated new vessel inside the tumour mass (Fig 2 N and Sup. Video 5). Interestingly, in both cancer models, we observed vessel co-option (Fig 2 J, T), with cancer cells beginning to migrate following the vessels as a dissemination route as described in the literature^2^. Thus, our method allows real-time visualization of tumor dissemination with the migration of cancer cells along the vessels. Simultaneously vessel morphology changes that promote tumor growth can be monitored. Real-time observation of vessel dynamics within the tumour is a valuable tool for the selection of new anti-tumour compounds such as anti-angiogenic treatments.

**Figure 2.**
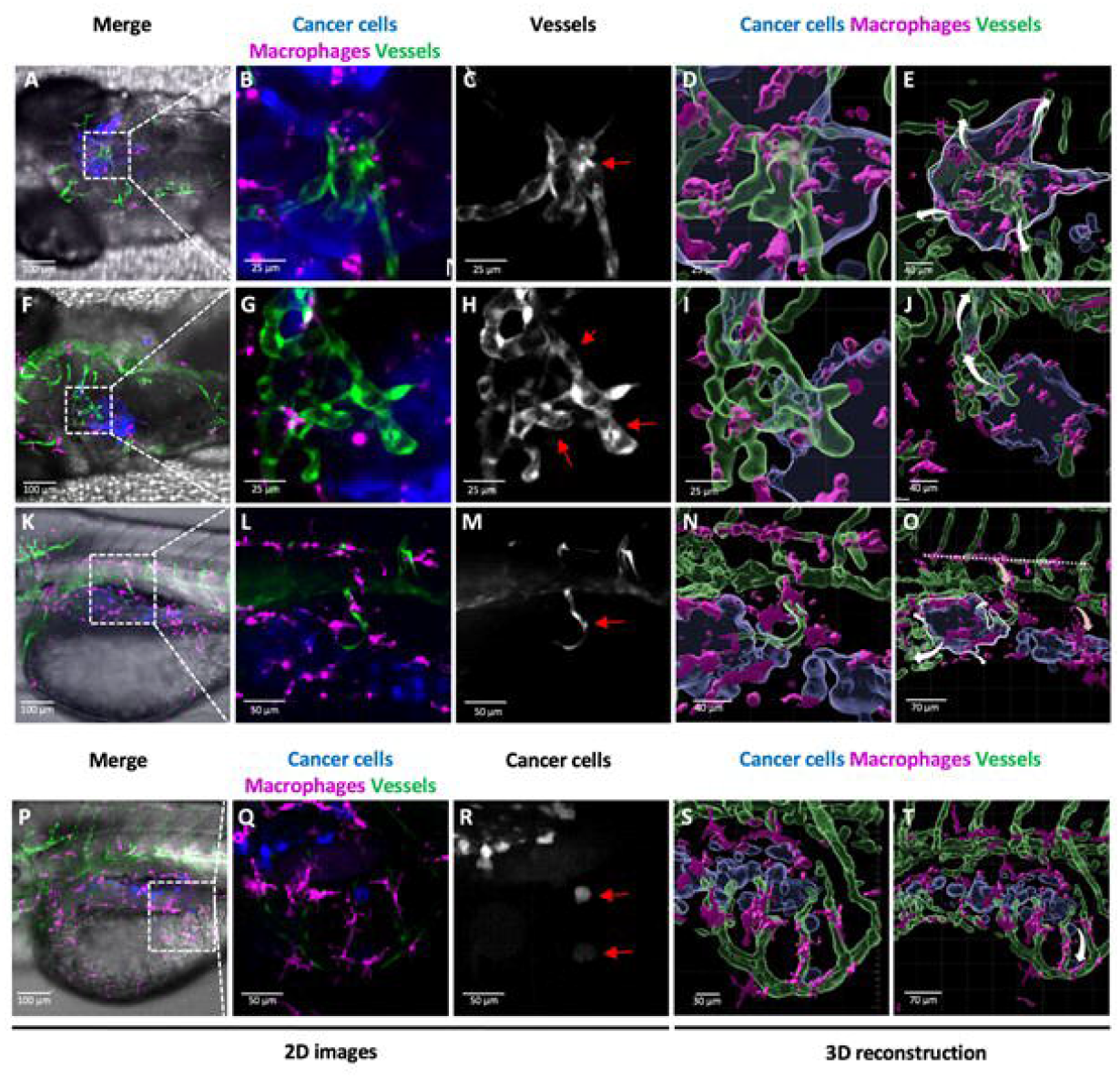
Analysis of 3D TME reconstruction and vessel dynamics: a predictive tool for tumor fate. **A-T**. Each cell population is highlighted with a coded color; cancer cells in blue or white, macrophages in magenta or white, and vessels in green or white as shown in the images. The arrows in 3D reconstructions represent the prediction of tumor fate (white arrow) and macrophage fate (light-red arrow). **A-E**. Z stack projection (A-C) and reconstruction (D-E) of the glioblastoma TME live imaging 48 hours after transplantation. **B-** **D**. The images show vessel dilatation in the center of the tumor mass (**C**, red arrow). **E**. The white arrow in the 3D reconstruction represents the prediction of tumor growth. **F-J**. Z stack projection (F-H) and reconstruction (I-J) of the glioblastoma TME live imaging 48 hours after transplantation. Images represent a Z projection (Z=30 slices) of the TME. **H**. Vessel angiogenesis in contact with the tumor mass (red arrow). **I-J**. 3D reconstruction showing cancer cells migration through the vessels in the upper left corner. **J**. The white arrow represents the predicted route of cancer dissemination regarding vessel localization. **K-O**. Z stack projection (K-M) and reconstruction (N-O) of the melanoma TME live imaging 24 hours after transplantation. Images represent a Z projection (Z=74 slices) of the TME. **L-M**. Neo-angiogenesis is shown with a new vessel redirected inside the tumor mass (red arrow). **N**. Macrophages are recruited at the tumor site following the new growing vessel. **O**. The horizontal dashed line represents macrophages orientation outside the tumor site. The light-red arrows represent the orientation of the macrophage migration. The white arrows represent the predicted route of cancer cells dissemination regarding vessel localization. **P-T**. Z stack projection (P-R) and reconstruction (S-T) of the melanoma TME live imaging 24 hours after transplantation. Images represent a Z projection (Z=74 slices) of the TME. **R**, In the images two cancer cells detach from the primary mass following the vessels (red arrows). **T**, The white arrow represents cancer cell migration and dissemination from primary mass.

Moreover, Figure 2 N-O shows the evolution of macrophage shape and orientations in space, from horizontal to vertical orientation, which allows their migration following the vasculature inside the tumour niche. Thus, considering the importance of macrophages in tumour angiogenesis^19^ the 3D reconstruction gives an insight of the crosstalk between these immune cells and vessels in the promotion of tumour progression.

Here we demonstrate the feasibility of modelling in 3D the dynamics of the TME at cellular and tissular level. Thanks to our methods, we pave the way for a complex understanding of the TME architecture modification *in vivo* by visualising the co-operation of specific cellular types such as macrophages and vessels, during the tumour progression. This method can be implemented by using other zebrafish transgenic lines to study other TME players such as neurons ^20^. At the same time, this tool can be applied to study other cancer types according to the different human cell lines used and the injection site. In this perspective, the 3D modelling of Patients Derived Xenografts (PDXs) in zebrafish embryos, would help clinicians in the treatment decision. Combining in the same model different colours allow to study the involvement of different cellular types in one embryo, saving time by performing less experiments thus decreasing experimental procedures cost. This is in line with the increased attention on the ethical guidelines. Finally with the exponential interest in precision medicines, this time-efficient and reliable method, consisting in 3D modelling of TME in zebrafish embryos, will represent a powerful tool to predict tumour fate in patients and to screen new efficient therapies.

## Acknowledgements

We thank V. Diakou and E. Jublanc from MRI platform for their help during image acquisition and image analysis. This work was supported by Agence Nationale de Recherche Technologie (ANRT), NEUcrest-ITN Project (EU’s Horizon 2020 grant No. 860635) and AZELEAD.

## Author

JM, FL, LF designed the experiments. JM, FL performed the experiments. JM, FL, KK and LF wrote the manuscript.

## Supplementary videos

**Supplementary Video 1: Navigation inside a 3D glioblastoma reconstruction to visualize macrophage recruitment and vessel organization**.

U87 human glioblastoma cells (blue) and its microenvironment such as macrophages (magenta) and vessels (green) are observed in the video. Human glioblastoma cells are transplanted in the midbrain region of zebrafish embryo at 3dpf, the video represents the glioblastoma TME 24h post transplantation. Macrophages are recruited all around the tumour mass and display close contact with glioblastoma cells suggesting an intensive communication between both populations.

**Supplementary Video 2: Navigation inside a 3D melanoma reconstruction to visualize macrophage recruitment**.

Human A375 melanoma cells (blue), macrophages (magenta) and vessels (green) are observed in the video. Human melanoma cells are transplanted in the swim bladder of a zebrafish embryo at 3dpf. The video represents the melanoma TME at 24h post transplantation. The navigation inside the TME 3D reconstruction allows to visualize the massive presence of macrophages around and inside the tumour mass.

**Supplementary Video 3: Macrophage dynamics in the glioblastoma tumour microenvironment**.

4D time-lapse reconstruction of U87 glioblastoma cells (blue) and macrophages (magenta). Human glioblastoma cells are transplanted in the midbrain region of zebrafish embryo at 3dpf, the video represents the glioblastoma TME at 24h post transplantation. The time Interval between two images is 7 min. Two macrophages behavioural dynamics is highlighted. Macrophages in contact with glioblastoma cells (white arrowhead) tends to have a low motility profile that allows long lasting contact between the two populations. On the opposite, peripheral macrophages (yellow arrowhead), distant from the tumor are more dynamic.

**Supplementary Video 4: Glioblastoma TME dynamics**.

4D time-lapse reconstruction of U87 glioblastoma cells (blue) and its microenvironment such as macrophages (magenta) and vessels (green). Human glioblastoma cells are transplanted in the midbrain region of a zebrafish embryo at 3dpf, the video represents the glioblastoma TME 24h post transplantation. The time Interval between two images is 7 min. The reconstruction allows the visualisation of vessel (white arrow) inside the core of tumour mass. Image acquisition allows to visualise at the same time multiple events such as mitosis (yellow asterisk *) and macrophage migration along vessels (white arrowhead). The spatial position of the cancer mass in the TME is fundamental to understand the mechanistic role of vessels.

**Supplementary Video 5: Navigation inside a 3D melanoma reconstruction to visualize vessel dynamics**.

Human A375 melanoma cells (blue), macrophages (magenta) and vessels (green) are observed in the video. Human melanoma cells are transplanted in the swim bladder of a zebrafish embryo at 3dpf, the video represents the melanoma TME 24h post transplantation.

The navigation inside the 3D reconstruction allows to visualize the dynamics of neo-angiogenic events. The spatial position of the cancer mass in the TME is fundamental to understand the mechanistic role of vessels. The video highlights the presence of a neo-growing-vessel inside the tumour mass located in the swim bladder where normally there is no vascular system. The specific localization of the vessel inside the tumour using the 3D reconstruction permit to predict the growth of the tumour itself.

## Online Material and Methods

### Animal rearing

Adult zebrafish (*Danio rerio*) were maintained on a 12/12 h light/dark cycle in a partially recirculating system and fed 3 times/day with fresh Artemia salina and dry food. Outcross of homozygous transgenic zebrafish lines *tg(KDR:eGFP)* and *tg(mfap4:RFP)* were used to perform cell transplantations. All experimental procedures on zebrafish were performed in accordance with the European guidelines and regulations on Animal Protection from the French Ministry of Health (F341725).

### Cell culture maintenance

Human glioblastoma cancer cell line, U87 WT, was purchased at ATCC (U-87 MG ATCC® HTB-14™). Human melanoma cancer cell line, A375 WT, was kindly provided by A. Pelegrin from IRCM - Montpellier. U87 and A375 cells were cultured in DMEM (Eurobio scientific) supplemented with 1% L-glutamine 200mM (Eurobio scientific), 1% Penicillin/Streptomycin (Eurobioscientific #CABPES01-0U) and 10% foetal bovine serum (Eurobio scientific) at standard conditions of 5% CO2, at 37°C. Cells were cultured in 100mm Petri dish (Corning) and split twice a week when they reached 80-90% confluency.

### Generation of stable fluorescent human cancer cell lines

To obtain U87-eBFP+ (blue fluorescent protein) and A375-eBFP+ cells, U87 WT and A375 cells were stably transfected with EBFP2-N1 plasmid using Phosphate calcium. EBFP2-N1 was a gift from Michael Davidson (Addgene plasmid #54595). One day before transfection, cells were plated in 6 well plates. EBFP2-N1 purified plasmid (4µg) was transfected using 2.5M CaCl2 reagent. 24h after transfection, complete DMEM medium was replaced. eBFP+ cells were selected using Geneticin (800-1000µg/mL) diluted in DMEM media during 4-6 weeks before starting *in vivo* experiments.

### Embryo and cell preparation prior to xenotransplantation

Embryos were initially maintained at 28 °C at a maximum density of 50 embryos per Petri dish in fish water supplemented with 0.0002% methylene blue (Sigma) as an antifungal agent. After 24h, embryos were placed in fish water containing 200µM Phenylthiourea (PTU) to prevent embryo pigmentation and allow fluorescent imaging acquisition.

Glioblastoma cells (U87-eBFP+) and melanoma cells (A375-eBFP+) were harvested from a 100mm petri dish the day of xenograft transplantation. Cells were washed with phosphate-buffered saline (PBS) and detached using Trypsine-Versene EDTA (Eurobio scientific) at 37°C for 5 min. Trypsine-versene was inactivated by complete DMEM media and cells were pelleted (1000rpm-5min) then washed with PBS. Finally, cells were resuspended in 50µL PBS to ensure highly concentrated cell preparation.

### Xenotransplantation of human cancer cell lines

Borosilicate glass capillaries (O.D.: 1 mm, I.D.: 0,75 mm, Sutter Instrument) were pulled using glass micropipette puller (Sutter Instrument). Capillaries were filled with 10 µL of cells suspension. Injection was performed under a stereomicroscope (Leica M80 Stereo zoom microscope) using a micromanipulator (Narishige) and oil manual microinjector (Cell Tram Vario Eppendorf). Before xenografting, embryos were anesthetized in 0.16 mg/ml PBS/Tricaine (MS-222) then tidily aligned laterally for melanoma or dorsally for glioblastoma transplantation. Embryos were maintained in position during injection using forceps. Melanoma cells were transplanted in the swim bladder, while glioblastoma cells were transplanted in the midbrain region. Between 50 to 100 cells were transplanted per embryo. Transplanted embryos were maintained at 33°C in fish water containing PTU.

### 3D and 4D live-imaging acquisition

A fluorescence stereomicroscope (AXIO Zoom.V16 Zeiss) equipped with eBFP filter was used for phenotypic selection of xenotransplanted embryos. Embryos with fluorescent tumor mass in the swim bladder or in the brain were selected for image/time lapse acquisition. 4 or 5 dpf (1- or 2-days post transplantation) anaesthetized embryos were mounted laterally or dorsally in 0,7% low melting point agarose (Life Technologies) in a 35mm fluorodish covered with fish water containing Tricaine. Images and time-lapse were acquired at 33°C using a Confocal spinning disk ANDOR coupled to a Nikon Ti Eclipse microscope (Objectives: 20x/0.75). 405/488/561 lasers were used with appropriate filters. For time-lapse, images were taken each 7 minutes with 1µm z-step interval.

### Image analysis

Images and time lapse were processed using Fiji and Imaris software. Images were smoothened and brightness/contrast was adjusted using Fiji software over all z-stacks. Maximum projections were generated using Fiji software. Segmentation and 3D reconstructions of macrophages, vessels and cancer cells were performed using Imaris Software 9.5. Supplementary videos were recorded using the same software.

